# Architecture of the Tuberous Sclerosis Protein Complex

**DOI:** 10.1101/2020.09.29.319707

**Authors:** Kailash Ramlaul, Wencheng Fu, Hua Li, Natàlia de Martin Garrido, Lin He, Wei Cui, Christopher H S Aylett, Geng Wu

## Abstract

The Tuberous Sclerosis Complex (TSC) protein complex (TSCC), comprising TSC1, TSC2, and TBC1D7, is widely recognised as a key integration hub for cell growth and intracellular stress signals upstream of the mammalian target of rapamycin complex 1 (mTORC1). The TSCC negatively regulates mTORC1 by acting as a GTPase-activating protein (GAP) towards the small GTPase Rheb. Both human TSC1 and TSC2 are important tumour suppressors, and mutations in them underlie the disease tuberous sclerosis.

We used single-particle cryo-EM to reveal the organisation and architecture of the complete human TSCC. We show that TSCC forms an elongated scorpion-like structure, consisting of a central “body”, with a “pincer” and a “tail” at the respective ends. The “body” is composed of a flexible TSC2 HEAT repeat dimer, along the inner surface of which runs the TSC1 coiled-coil backbone, breaking the symmetry of the dimer. Each end of the body is structurally distinct, representing the N- and C-termini of TSC1; a “pincer” is formed by the highly flexible N-terminal TSC1 core domains and a barbed “tail” makes up the TSC1 coiled-coil-TBC1D7 junction. The TSC2 GAP domain is found abutting the centre of the body on each side of the dimerisation interface, poised to bind a pair of Rheb molecules at a similar separation to the pair in activated mTORC1.

Our architectural dissection reveals the mode of association and topology of the complex, casts light on the recruitment of Rheb to the TSCC, and also hints at functional higher order oligomerisation, which has previously been predicted to be important for Rheb-signalling suppression.

## Introduction

Tuberous sclerosis complex (TSC) is an autosomal dominant disease characterised by benign tumours in multiple organs (Henske *et al*, 2016). It is caused by mutations in either of the genes *TSC1* or *TSC2*, which encode the 130 kDa TSC1 and the 200 kDa TSC2 tumour suppressor proteins respectively. TSC1 contains an N-terminal α-helical ‘core’ domain and a coiled-coil at the C-terminus which is required for binding TSC2 (Nellist *et al*, 1999; Santiago Lima *et al*, 2014; van Slegtenhorst *et al*, 1998). TSC2 contains a long α-solenoid domain at the N-terminus and a C-terminal GTPase activating protein (GAP) domain, which is the sole catalytically active domain in the complex. Together with a small third subunit TBC1D7 (Dibble *et al*, 2012), TSC1 and TSC2 assemble to form the TSC protein complex (TSCC).

TSCC signalling restricts cell growth by negatively regulating mTORC1, the central coordinator of metabolism (González & Hall, 2017; Ramlaul & Aylett, 2018). Directly upstream of mTORC1, Rheb, a small GTPase localized to lysosomes through C-terminal farnesylation (Clark *et al*, 1997), stimulates mTORC1 kinase activity when GTP-bound (Yang *et al*, 2017). The TSCC stimulates Rheb GTPase activity, accumulating the GTPase in the inactive, GDP-bound, state to suppress mTORC1 (Huang & Manning, 2008). Spatial regulation of TSCC between the cytoplasm and lysosome is known to be pivotal for its function as a Rheb-GAP, with the current understanding being that the TSCC translocates to the lysosome surface to catalytically and sterically inhibit mTORC1 by binding to Rheb and sequestering it (Menon *et al*, 2014; Demetriades *et al*, 2016). This translocation is reversed on TSC2 phosphorylation by AKT (Menon *et al*, 2014), and other kinases, which are thought to regulate localisation through an unknown mechanism involving 14-3-3 binding (Shumway *et al*, 2003; Cai *et al*, 2006), as well as TSCC breakdown by ubiquitination-targeted TSC2 degradation (Benvenuto *et al*, 2000; Chong-Kopera *et al*, 2006).

The architecture of the TSCC remains completely unknown, although small fragments of the complex have been structurally characterised. The core domain of *S. pombe* TSC1 (Sun *et al*, 2013), an N-terminal α-solenoid fragment of *C. thermophilum* TSC2 (Zech *et al*, 2016), and most recently the *Ct*TSC2GAP domain (Hansmann *et al*, 2020) have been resolved crystallographically. Furthermore, two co-crystal structures of TBC1D7 interacting with C-terminal coiled-coil fragments of TSC1 have been determined (Gai *et al*, 2016; Qin *et al*, 2016). In this study, we have used cryogenic electron microscopy (cryo-EM) to examine the molecular architecture of the full-length, human holo-TSCC.

## Results

We cloned human TSC1, TSC2, and TBC1D7 for expression in human embryonic kidney cells, and Rheb for expression in *Escherichia coli*. A clone yielding an internal deletion of TSC1(Δ400-600) was used for SEC-MALLS to minimise inter-complex interactions. While both TBC1D7 and TSC1 could be expressed and purified independently, TSC2 could not be purified in the absence of TSC1 (Supp. Fig. 1A), forming inclusions or being degraded in cell, consistent with the role of TSC1 in preventing TSC2 degradation (Benvenuto *et al*, 2000; Chong-Kopera *et al*, 2006).

We retrieved the complete human TSCC from lysate using FLAG-tagging, and purified the TSCC from most remaining contaminants by size-exclusion chromatography (SEC) (Supp. Fig. 1B). Purified TSCC exhibited its physiological GAP activity towards Rheb (Supp. Fig. 1C). Purification and SEC of TSC1(Δ400-600) TSCC yielded a defined peak with an estimated molecular weight by multi-angle LASER light scattering of 700 kDa (Supp. Fig. 1D), roughly corresponding to a composition of 2:2:1-TSC1:TSC2:TBC1D7. Comparatively, the full-length TSCC yielded a broad peak with an estimated mass of 5,200 kDa, due to lateral inter-TSCC interactions (Supp. Fig. 2A).

We initially investigated the molecular architecture of the TSCC by negative staining. We observed extremely flexible, independent particles, however three defined ordered regions could be isolated, and two-dimensional averaging of these regions provided a complete picture (Fig. 1A). The TSCC was extraordinarily elongated (~400 Å) and exhibited a characteristic “scorpion” shape, with a bulkier central “body”, flexible “pincer”-like appendage at one end, and a barbed “tail” at the other. Cryo-EM of TSCC at low concentrations revealed identical particles, whereas at higher concentrations we observed web-like networks which appear to be formed of head-to-tail TSCC particles (Supp. Fig. 2B). Once again, complete TSCC particles were too flexible to average beyond low resolution. We isolated the same regions; both the “pincer” and “tail” proved to be strongly preferentially oriented and flexible, refining only to low-intermediate resolution (8.1 Å and 8.2 Å respectively) (Fig. 1B, Supp. Fig. 3). The body of the TSCC exhibited pseudo-C2 symmetry, and was refined in C2 initially, before symmetry was relaxed for a final C1 structure at 4.6 Å (Fig. 1B, Supp. Fig. 3).

**Figure 1:**
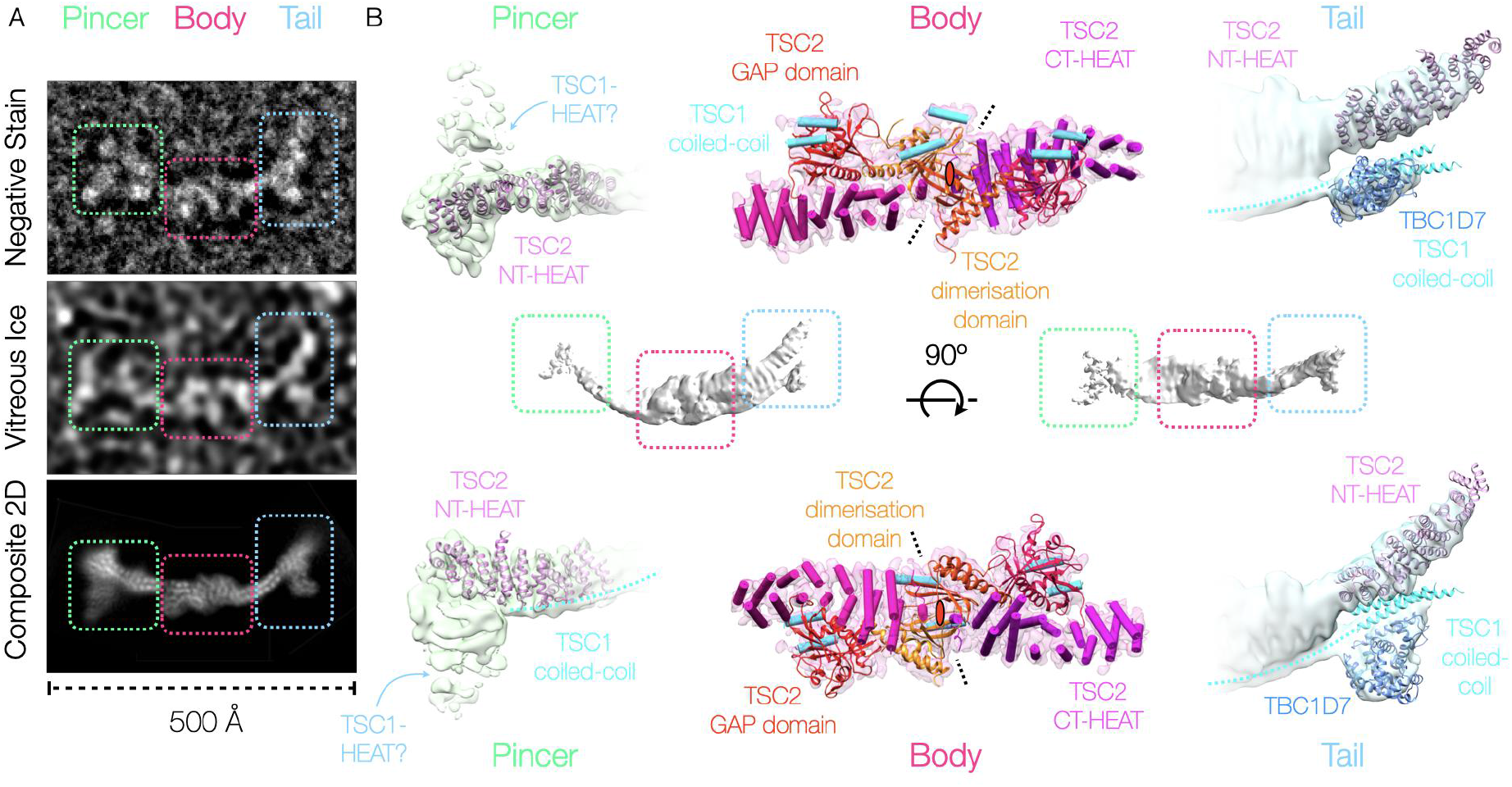
The TSC protein complex is an elongated, flexible, scorpion-like complex with a defined “pincer”, “body”, and barbed “tail”. [A] Electron micrograph of a negatively stained TSCC particle on a carbon support, electron micrograph of a TSCC particle frozen within vitreous ice on a graphene oxide support, and composite 2D average image of the TSCC from the windowed regions of vitrified particles as indicated. The same regions, “pincer” (chartreuse), “body” (cerise), and “tail” (cerulean), are indicated through dashed boxes of the appropriate colour in both the representative particle image and the composite 2D class-average representation. [B] The overall structure of TSCC at low resolution (centre) and the refined densities corresponding to each region of the TSCC (indicated by boxes) are shown. In each case the reconstructed electron scattering density is shown as a transparent isosurface, while the corresponding fitted molecular structures (the TSC2 N-terminal HEAT-repeat (Zech *et al*, 2016), the TBC1D7-TSC1 complex (Gai *et al*, 2016; Qin *et al*, 2016), the TSC2-GAP domain (Hansmann *et al*, 2020), and the RapGAP dimerisation domain (Scrima *et al*, 2008)), and secondary structural elements in the case of the body, are shown in cartoon representation where available and practicable. The reconstructions have been rotated by 90° in the second panel as indicated.

The resolution of the “body” was high enough to trace chains and identify all secondary structural elements, but too low to definitively assign sequence. The published structure of the TSC2 GAP domain (Hansmann *et al*, 2020) fitted unambiguously into density adjoining a central α-solenoid on each side of the origin of C2 pseudo-symmetry (Supp. Fig. 4A). At the juncture of the two α-solenoids, we observed a dimerisation interface comprising two back-to-back β-sheets. The topology of this dimerisation domain is conserved from that of the N-terminal domain of the RapGAP fold (Scrima *et al*, 2008) (Supp. Fig. 4B), although a domain swap between the two TSC2 molecules cannot be ruled out as the regions of secondary structure are separated by hundreds of disordered residues (Supp. Fig. 5). With the exception of the GAP domain, the only β-elements remaining predicted within the sequence of any TSC protein are at the C-terminus of the α-solenoid of TSC2 (Supp. Fig. 5), implying that the long disordered regions containing many of the phosphorylation sites regulating the TSCC are insertions within the C-terminal GAP domain. Our results are consistent with the TSC2 α-solenoids running outwards from C-terminus to N-terminus from the dimerisation site, and indeed there is a good fit of the TSC2 N-terminal HEAT repeat structure into the end of each of the “pincer” and “tail” (Fig. 1B, Supp. Fig. 4C/D). The C2 symmetry of the TSC2 dimer is broken by two helices running directly across the top of the RapGAP-like dimerisation domain. This helical density forms a weakly connected “backbone” running over both GAP domains, and along the TSC2 α-solenoid outwards to both the “pincer” and “tail”. We assign this continuous helical coiled-coil as that from the C-terminal regions of TSC1, implying that the two ends are its N- and C-terminus respectively. The “pincer” density is uninterpretable, however the TSC1-TBC1D7 structure (Qin *et al*, 2016) unambiguously fits the density corresponding to the “barb” lying on the α-solenoid of the “tail” (Supp. Fig. 4D), confirming the orientation of the TSC1 dimer, and implying that the “pincer” is made up of the TSC1 HEAT-domains.

Under the reasonable assumption that the GAP-Rheb interaction will mirror that of the published Rap-RapGAP complex (Scrima *et al*, 2008), we have docked Rheb accordingly (Supp. Fig. 6A). The natural docking yields no clashes with the current structure, and implies a further interaction with the two helices adjoining the GAP domain which are conserved from the RapGAP fold (Fig. 2B). We note that the Rheb farnesylation sites would both be situated on the same side of the TSCC, consistent with this being the correct orientation for lysosomal binding.

**Figure 2:**
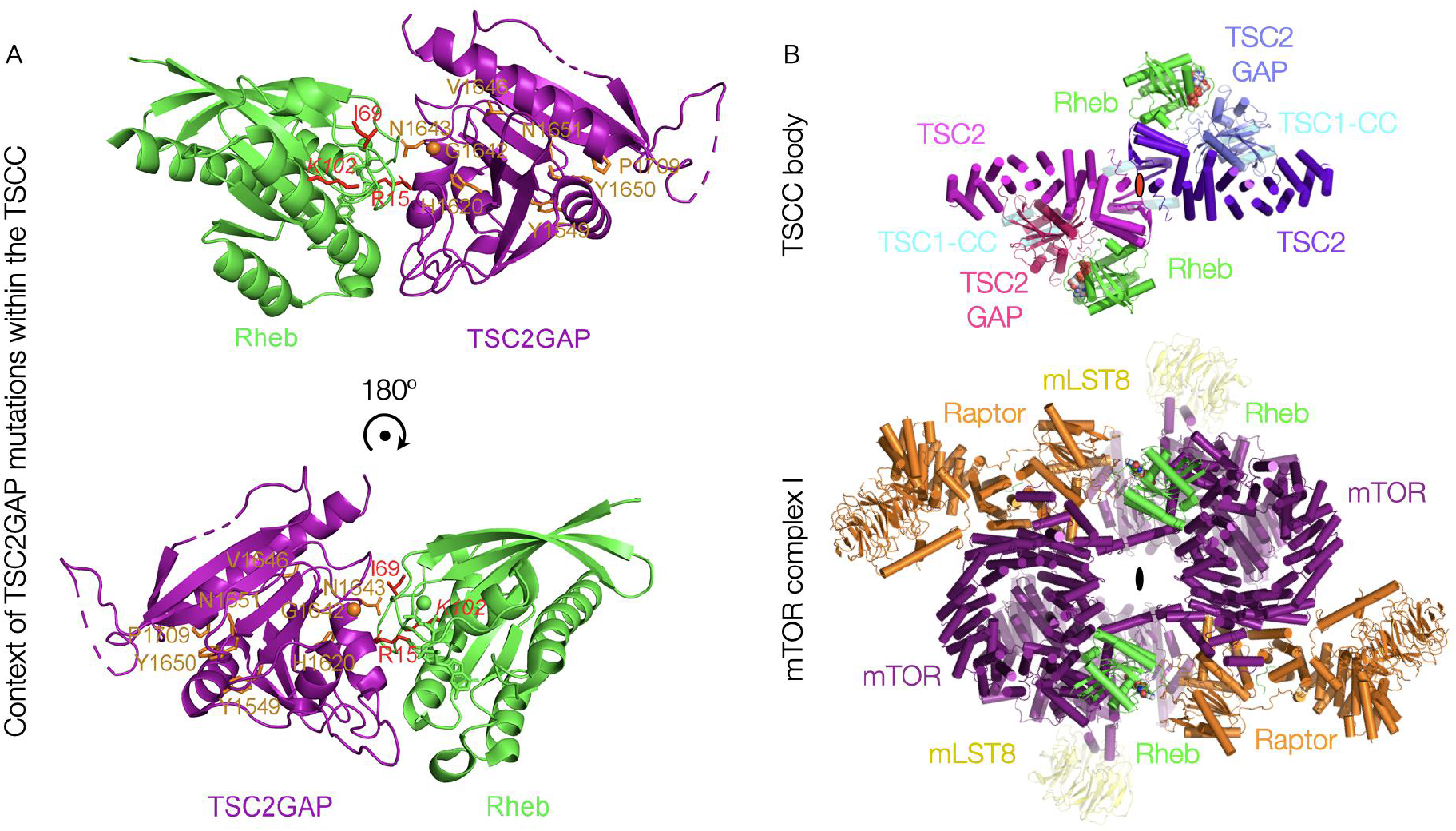
The TSCC central TSC2 GAP domains are poised to bind two Rheb molecules in interactions expected to be disrupted by key tumorigenic mutations. [A] A model of TSC2-GAP in complex with Rheb. The crystal structures of *Ct*TSC2_GAP_ domain and human Rheb are superimposed onto the crystal structure of the RapGAP-Rap complex. *Ct*SC2_GAP_ residues corresponding to human TSC2 residues targeted by tumorigenic mutations in tuberous sclerosis are shown in stick representation and coloured in orange, labelled with human TSC2 residue numbers. The three Rheb residues identified to be conserved in Rheb homologues and Rheb-specific among Ras family GTPases are displayed in stick representation and coloured in red. The second panel has been rotated by 180° as indicated. [B] The docked fit of Rheb against the TSC2 GAP domain, based on the structure of the RapGAP-Rap1 complex, within the “body” of the TSCC, in comparison to its fit in the Rheb-activated structure of mTORC1. The secondary structural elements of the TSCC, and the molecular structure of mTORC1, are shown in cartoon representation.

## Discussion

We show that the TSCC forms an elongated, flexible architecture, comprising two copies of each of TSC1 and TSC2 and one of TBC1D7. The orientation of Rheb implied by the GAP domains (Fig. 2B) matches the slight curvature of the complex, and the lysosomal membrane will therefore lie on the opposite side of the TSCC from the TSC1 backbone. The super-structure observed forming at higher concentrations (Supp. Fig. 2B) may well play a part in retaining TSCC at the lysosome and reducing the off-rate once Rheb signalling has been suppressed as previously predicted (Menon *et al*, 2014). Further structural investigation of these inter-TSCC interactions, likely mediated by the TSC1 termini (Nellist *et al*, 1999; Hoogeveen-Westerveld *et al*, 2010), is required to understand the higher-order organisation of TSCC and its role in mTORC1 regulation.

The RapGAP-like domain of TSC2 forms a dimer, as reported by Scrima and colleagues (Scrima *et al*, 2008), providing the centre of pseudosymmetry of the TSCC. The C2 symmetry of each of the dimeric TSC proteins is broken by the presence of the other, that of TSC1 by the curvature of its coiled-coil along the TSC2 α-solenoids, and that of TSC2 by the involvement of TSC1 in its dimerisation. We have confirmed once again that while TSC1 can fold independently of TSC2, the reverse does not occur (Benvenuto *et al*, 2000; Chong-Kopera *et al*, 2006; Woodford *et al*, 2017). Our architecture suggests a structural explanation for this observation; direct TSC1 involvement in the TSC2 dimerisation interface. The previously observed breakdown of TSC2, following ubiquitination in the absence of TSC1 (Benvenuto *et al*, 2000; Chong-Kopera *et al*, 2006), would be expected when structural elements cannot fold appropriately in the absence of their partner. This would also constrain the presence of functional TSC2 to subcellular regions containing TSC1 dimers.

*tsc2* mutations are highly correlated with tuberous sclerosis. Half of these tumorigenic missense mutations occur within the N-terminal domain (NTD) of TSC2 (27 of 54 residues in Supp. Table 1). Interestingly, most of these disease-targeting residues cluster together, such as A84/P91/E92, C244/M286/G294/E337/A357/R367, G440/L448/A460/R462/L466/L493, A583/H597/Y598/A607/R611/R622/M649, and L826/L847/R905/L916 (Supp. Fig. 5). Given that the TSC2-NTD closely contacts the TSC2-GAP and TSC1-coiled coil in our structure, these clusters within the TSC2-NTD may represent the GAP- or TSC1-interacting surfaces. The TSC2-GAP domain possesses the second most tumorigenic missense mutations, with >30% of all sites (17 of 54 in Supp. Table 1). Many disease-related residues are highly conserved, such as H1620/G1642/N1643/R1743 (Supp. Fig. 5). Residues H1620/G1642/N1643 in TSC2-GAP correspond to H266/G289/N290 of RapGAP, which directly contact or are very close to Rap in the Rap-RapGAP structure (Scrima et al., 2008, Fig. 2A, Supp. Fig. 6). These residues are expected to interact with Rheb. In cancer cells harbouring these *tsc2* mutations, the TSC2-NTD/TSC2-GAP, or TSC1/TSC2, or TSC2-GAP/Rheb associations may be disrupted, leading to diminished GAP activity toward Rheb and abnormally elevated mTORC1 activity.

In our structure, the N-terminal part of TSC1 coiled coil is found to interact with the N-terminal HEAT domain of TSC2. Interestingly, this region of TSC1 is the highest conserved part among TSC1 homologues (Supp. Fig. 7).

By comparing the sequences of Rheb homologues and other Ras family GTPases, we found three Rheb-specific residues: R15, I69, and K102. They are conserved in Rheb homologues but are different in all other Ras family GTPases (Supp. Fig. 8). Interestingly, these Rheb-unique residues all point toward TSC2-GAP in our model (Fig. 2, Supp. Fig. 6)

In complete agreement with the conclusions of the Manning group (Menon *et al*, 2014), the presence of the TSCC will both catalytically and sterically prevent Rheb–mTORC1 interactions (Fig. 2B), rotating the Rheb pair in relation to its position when interacting with mTORC1. We believe that one of the more interesting observations from our results is that TSC2 binds Rheb as such a pair, as does mTORC1. Despite the fact that they are completely different in architecture and approach from different directions, the TSCC GAP domains are poised to bind two copies of Rheb at an almost identical separation to that resolved for Rheb in the structure of activated mTORC1 (Fig. 2B). While it is possible that this is entirely happenstance, this would also be expected were Rheb bound by each partner as part of a greater, at least dimeric, complex on the lysosomal surface.

Our improved architectural understanding of the TSCC provides a starting point for the investigation of the molecular mechanisms by which TSCC directly regulates Rheb, and poses new questions on the nature of the superstructures formed by TSCC complexes, their partners, and the involvement of such quaternary structures in mTORC1 regulation.

## Materials and methods

### Protein expression and purification

pRK7 plasmids subcloned with FLAG-tagged full-length (FL) human TSC1 (1164 amino acids, UniProtKB/Swiss-Prot accession number Q92574-1) and FLAG-tagged FL human TSC2 (1807 amino acids, UniProtKB/Swiss-Prot accession number P49815-1) were purchased from Addgene, and pRK7 was subcloned with FLAG-tagged human TBC1D7 (293 amino acids, GenBank accession number AAH07054). FL TSC1-TSC2-TBC1D7 (TSCC_FL_) plasmids, or TSC1_Δ400-600_-TSC2-TBC1D7 (TSCC_1Δ_) plasmids, were co-transfected into human embryonic kidney (HEK) Expi293F cells (Thermo Fisher Scientific, Waltham, MA, USA). Two days after transfection, the harvested Expi293F cells were lysed by three cycles of freeze-thaw in lysis buffer (20 mM Tris, pH 8.0, 300 mM NaCl, 2 mM TCEP, 0.5 mM PMSF, 1 μg/ml aprotinin, and 1 μg/ml leupeptin), and TSCC was purified from the cell lysate by M2 anti-FLAG affinity chromatography (Sigma) followed by size exclusion chromatography using a Superose 6 column (GE Healthcare) pre-equilibrated with buffer containing 20 mM Tris, pH 8.0, 300 mM NaCl, and 2 mM TCEP. The identity of each TSCC component was verified by ESI-MS (Mass Spectrometry Facility, University of St. Andrews).

The cDNA residues 1-169 of human Rheb (184 amino acid isoform, GenBank accession number EAW53989) was subcloned into the pET28a vector (Novagen), with an N-terminal 6×His-tag. Rhebwas overexpressed in *E.coli* strain BL21(DE3). After lysis of the bacteria with a cell homogenizer (JNBIO) and clarification, the lysate was subjected to Ni^2+^-NTA affinity chromatography (Qiagen) followed by size exclusion chromatography using a Superdex 200 column (GE Healthcare) pre-equilibrated with buffer containing 20 mM Tris, pH 8.0, 200 mM NaCl, and 2 mM DTT.

### GTPase activity endpoint assay

The GTPase activity of Rheb was assayed using the QuantiChrom ATPase/GTPase assay kit (BioAssay Systems), in which the amount of released inorganic phosphate was measured through a chromogenic reaction with malachite green. In the assays 75 nM RheB, either alone or mixed with 227.5 nM TSCC_FL_, were added to the reaction buffer (40 mM Tris, pH 8.0, 80 mM NaCl, 8 mM MgOAc, 1 mM EDTA and 14 mM GTP) at 28 °C for 40 min. A further 200 μL of assay kit reagent was then added, and the reaction incubated for 20 min, before a microplate reading at OD 620 nm was measured. Spontaneous GTP hydrolysis was calculated by measuring background absorbance in the absence of Rheb and sample valueswere normalised by subtraction of background. Each experiment was repeated three times.

### Size exclusion chromatography–multi-angle laser light scattering (SEC-MALLS)

TSCC_1Δ_ was analysed by SEC-MALLS using an Infinity liquid chromatography system (Agilent Technologies), linked to a Dawn Heleos multi-angle light scattering detector (Wyatt Technology) and Optilab T-rEX refractive index detector (Wyatt Technology). The sample was injected onto a Superose 6 10/300 size exclusion column (GE Healthcare) pre-equilibrated overnight with buffer containing 25 mM K-HEPES, pH 7.6, 250 mM KCl, 0.5 mM EDTA, 1 mM TCEP and trace amounts of NaN_3_, using 0.2 mL/min flow rate at room temperature. In-line UV absorbance, light scattering and refractive index measurements were analysed using the ASTRA software package (Wyatt Technology) to determine molar mass estimates.

### Sample preparation for cryo-EM studies

TSCC, after the above purification steps, was loaded onto a Superose 6 10/300 size exclusion chromatography column (GE Healthcare) pre-equilibrated with a preparation buffer containing 25 mM K·HEPES, pH 7.6, 175 mM KCl or 150 mM LiCl, 1 mM TCEP, and 0.5 μM EDTA. TSCC eluted as a single peak with a slight shoulder at lower retention volume. The integrity of the complex was confirmed by SDS-PAGE of both the peak and shoulder fractions. Main peak fractions were combined and concentrated to 0.1–0.2 mg/mL using Amicon 100 kDa molecular weight cut-off (MWCO) centrifugal filters and used for grid preparation.

### Generation of an initial TSCC reference density

A sample of concentrated wild-type TSCC_FL_ was applied to a carbon-coated holey carbon grid (R1.2/1.3, Quantifoil) and stained with 2% (w/v) uranyl acetate. A total of 224 micrographs were collected using an FEI Tecnai T12 electron microscope (Thermo Fisher Scientific, Waltham, MA, USA) at a magnification of 81,000-fold, an acceleration voltage of 120 kV, and a total dose of 50 e^-^ /Å^2^ over a defocus range of −0.5 to −2.0 μm. A dataset of 9597 particles was selected manually using BOXER. The parameters of the contrast transfer function were determined using CTFFIND4. Particles were 2D-classified into 100 classes in two dimensions using RELION and sixteen well-defined classes were selected for initial three-dimensional reconstruction. Initial models were created using the initial model functions in EMAN2, refined in three dimensions at low resolution using SPIDER, then filtered to 60 Å and used as an initial reference for automatic refinement in RELION. The resulting initial model at a resolution of 26 Å, with independent volume Fourier Shell Correlation (FSC) of 0.143, was used for further refinement.

### TSCC cryo-EM sample preparation

Samples of concentrated TSCC_FL_ were adsorbed to a thin film of graphene oxide deposited upon the surface of holey carbon copper grids (R2/1, 300 mesh, Quantifoil). Grids were blotted for 2-3 seconds before plunge freezing in liquid ethane using a Vitrobot Mark IV (Thermo Fisher Scientific, Waltham, MA, USA) at 4 °C and 100% humidity.

### TSCC cryo-EM data collection

Data was collected of TSCC on a Titan Krios (Thermo Fisher Scientific, Waltham, MA, USA) at the Electron Bioimaging Centre (eBIC, Diamond Light Source), equipped with a K2 Summit direct electron detector (GATAN, San Diego, USA) and operated at 300 kV, 37000-fold magnification and with an applied defocus range of −0.75 to −3.25 μm. Frames were recorded automatically using EPU, resulting in 5,387 images of 3,838 by 3,710 pixels with a pixel size of 1.35 Å on the object scale. Images were recorded in two successive datasets (of 1880 and 3507 images, respectively) as either 40 or 60 separate frames in electron counting mode, comprising a total exposure of 52.3 or 80.2 e^-^Å^−2^, respectively.

### TSCC cryo-EM data processing

Frames were aligned, summed and weighted by dose according to the method of Grant and Grigorieff using MotionCor2 (Zheng *et al*, 2017) to obtain a final image. Poor-quality micrographs were rejected based on diminished estimated maximum resolution on CTF estimation using CTFFIND4 (Rohou & Grigorieff, 2015) and visually based on irregularity of the observed Thon rings. Particles were selected using BATCHBOXER (Tang *et al*, 2007), and refinement thereafter performed using RELION_3_ (Scheres, 2012; Zivanov *et al*, 2018).

Two-dimensional reference-free alignment was performed on ~1,500,000 initially boxed particles to exclude those that did not yield high-resolution class averages and to identify the principal ordered regions of the TSCC molecule. Of these, 395,622particles populated classes extending to high-resolution and were retained for further refinement.

TSCC proved to be highly preferentially oriented on the grid, however it was possible to identify 2D classes for each of the “body”, “pincer” and “tail” regions. Iterated re-centring, twodimensional refinement, and re-boxing using the neural network particle picker Topaz (Bepler *et al*, 2018) was performed from the “body” region outwards in order to recover enough “pincer” and “tail” views to provide a complete description and definitive topology for all three regions.

Particles belonging to the “body” frequently displayed C2 symmetry in 2D class averages, and this particle subsetwas refined in three dimensions using this symmetry restraint. After several iterations of re-picking particles using Topaz (Bepler *et al*, 2018) and refinement, the final gold-standard refinement of the “body”, including 172,093 particles, reached 4.2Å at an independent FSC = 0.143. The symmetry was subsequently relaxed to C1 and refined (gold-standard) to 4.6 Å resolution at an independent FSC = 0.143. Reconstructions were also performed of the “pincer” and “tail” regions, from 15,854 and 58,307 particles respectively, however these suffered from persistent highly preferred orientation and conformational flexibility, with gold-standard refinements reaching 8.1 Å and 8.2 Å resolution, respectively, at an independent FSC = 0.143.

## Supporting information

Architecture of the Tuberous Sclerosis Protein Complex_supplementary

## Acknowledgements

The authors would like to thank Xinqiu Guo at Instrumental Analysis Centre of Shanghai Jiao Tong University for assistance with electron microscopy experiments. G.W. thanks Dr. Sheng Wang, Dr. Xinqi Gong, Lijun Yan, Hongpeng Wang, Ailiang He, and Dr. Yuan Gao for their contribution. We are indebted to the Imperial Centre for Structural Biology and the UK national electron Bioimaging Centre at Diamond Light Source (proposal EM14769, funded by the Wellcome Trust, MRC and BBSRC) for access to electron microscopy equipment and in particular to Paul Simpson for technical support. The authors would also like to thank Marc Morgan for help with SEC-MALLS analysis, the St. Andrews Proteomics facility for mass spectrometric protein identification and Nadezhda Aleksandrova and Zining Zhu for performing biochemical assays.

G.W. is supported by National Natural Science Foundation of China (grant numbers 31670106 and 31872627), Shanghai Jiao Tong University Scientific and Technological Innovation Fund, and the National Key R&D Program of China (YS2020YFA090044). C.H.S.A. is supported by a Sir Henry Dale Fellowship jointly funded by the Wellcome Trust and the Royal Society (206212/Z/17/Z).

## Author contributions

G.W. and C.H.S.A. conceived of the project. H.L., K.R., and W.F. purified the protein, performed initial negative staining analysis, and carried out the GAP assay. K.R. and N.M.G. carried out and analysed SEC-MALLS experiments. C.H.S.A., K.R., and W.F. prepared samples and grids for cryo-EM, collected electron micrographs and calculated single-particle reconstructions. All authors were involved in the interpretation of results and the drafting of the manuscript.

## Data and materials availability

The cryo-EM density maps corresponding to the “pincer”, “body”, and “tail” of the *HsTSCC* complex has been deposited in the EM Databank under accession codes EMD-11816, EMD-11819, and EMD-11817.

