## Supplementary material for "Architecture of the Tuberous Sclerosis Protein Complex": Architecture of the Tuberous Sclerosis Protein Complex_supplementary

### Supplementary Information

[1] Section for Structural Biology, Department of Medicine, Imperial College London, Exhibition Road, London, SW7 2BB, United Kingdom

[2] State Key Laboratory of Microbial Metabolism, School of Life Sciences & Biotechnology, The Joint International Research Laboratory of Metabolic & Developmental Sciences MOE, Shanghai Jiao Tong University, Shanghai, China

[3]Instrumental Analysis Center, Shanghai Jiao Tong University, Shanghai, China

[4]Institute of Reproductive and Developmental Biology, Faculty of Medicine, Imperial College London, Du Cane Road, London, W12 0NN, United Kingdom

* These authors contributed equally to this study.

† To whom correspondence may be addressed:

C.H.S.A. -

G.W. -

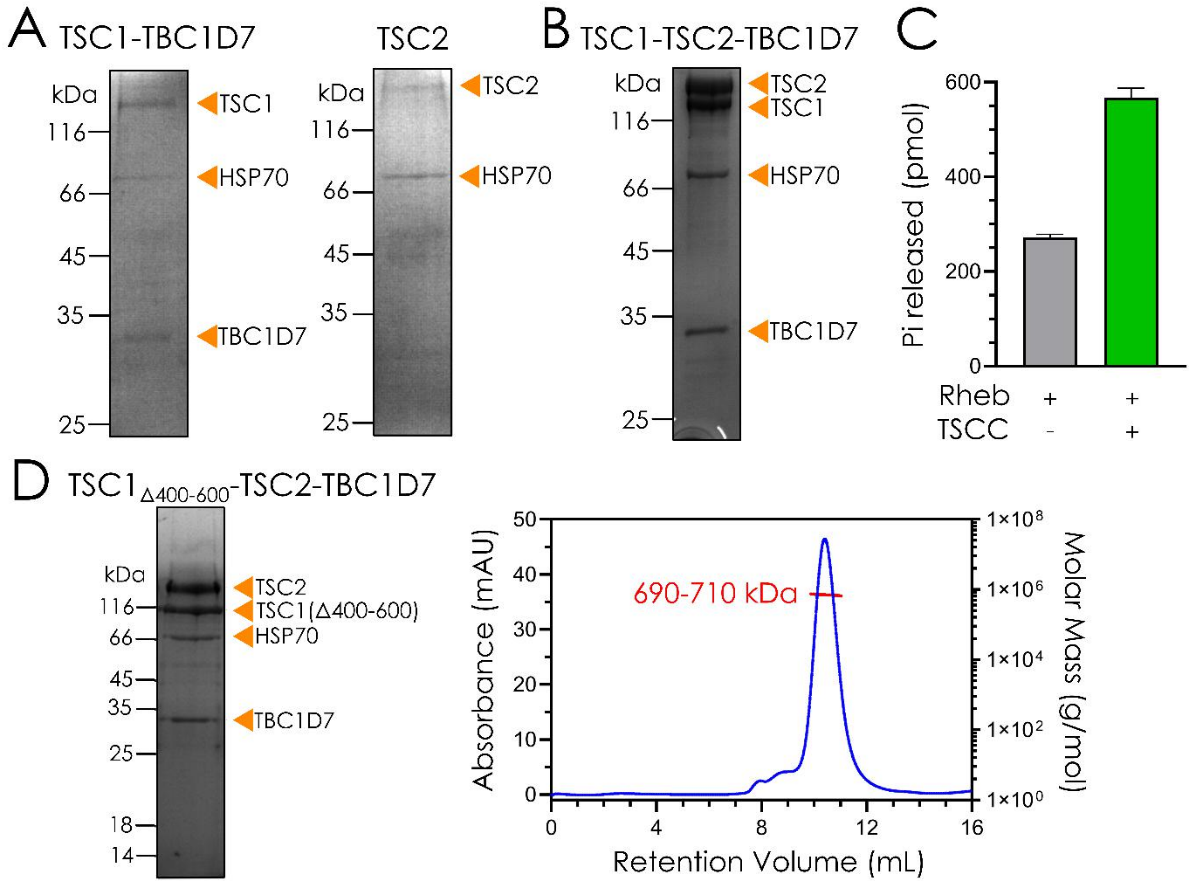


**Supplementary Figure 1. Production of functional human holo-TSCC.** A. Coomassie-stained SDS-PAGE gels showing TSC1-TBC1D7 alone (left) could be expressed and purified after transient transfection in HEK293F cells, whereas TSC2 alone (right) expressed in much lower quantity and was found to be irreversibly aggregated or degraded. B. Coomassie-stained SDS-PAGE of purified holo-TSCC, demonstrating stoichiometric pulldown of all three components in good yield. C. Results of three repetitions of a malachite green inorganic phosphate release assay conducted with purified Rheb and with purified Rheb and TSCC, showing a clear increase in activity in the presence of the TSCC. D. (Left) Coomassie-stained SDS-PAGE of purified $TSC1_{\Delta 400\text{-}600}$-TSC2-TBC1D7, indicating that deletion of residues 400-600 did not affect the integrity of the core complex. (Right) SEC-MALLS trace and molecular weight estimate for $TSC1_{\Delta 400\text{-}600}$-TSC2-TBC1D7. The peak was isotropic, and the molecular weight estimate was well within range of the observed protein components upon later structural examination.

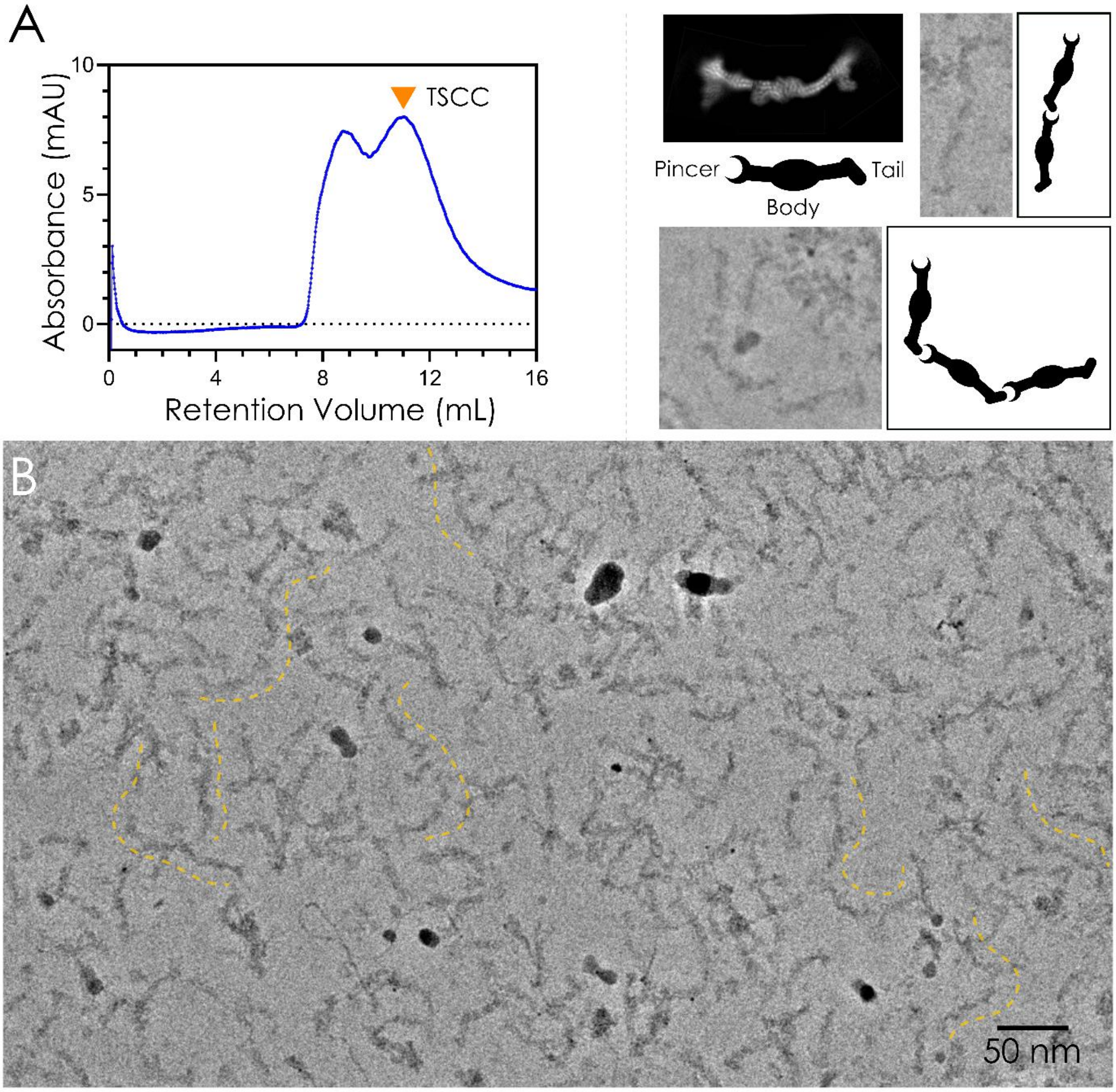


**Supplementary Figure 2. TSCC forms web-like networks, likely mediated by TSC1, which associate in pincer-to-tail arrangements.** A. Full-length TSCC elutes with an approximate molecular weight of 5200 kDa from a Superose 6 size exclusion column, indicating the presence of inter-TSCC interactions leading to higher-order oligomer formation. B. View of cryo-electron micrograph in which TSCC forms higher-order oligomers (orange dotted lines). These micrographs were collected on a Titan Krios fitted with a Volta phase plate for increased contrast. (Above, inset) Close comparison of TSCC oligomer views with our proposed architectural model of TSCC reveals that the complex forms these filaments with a defined, “pincer-to-tail”, topology.

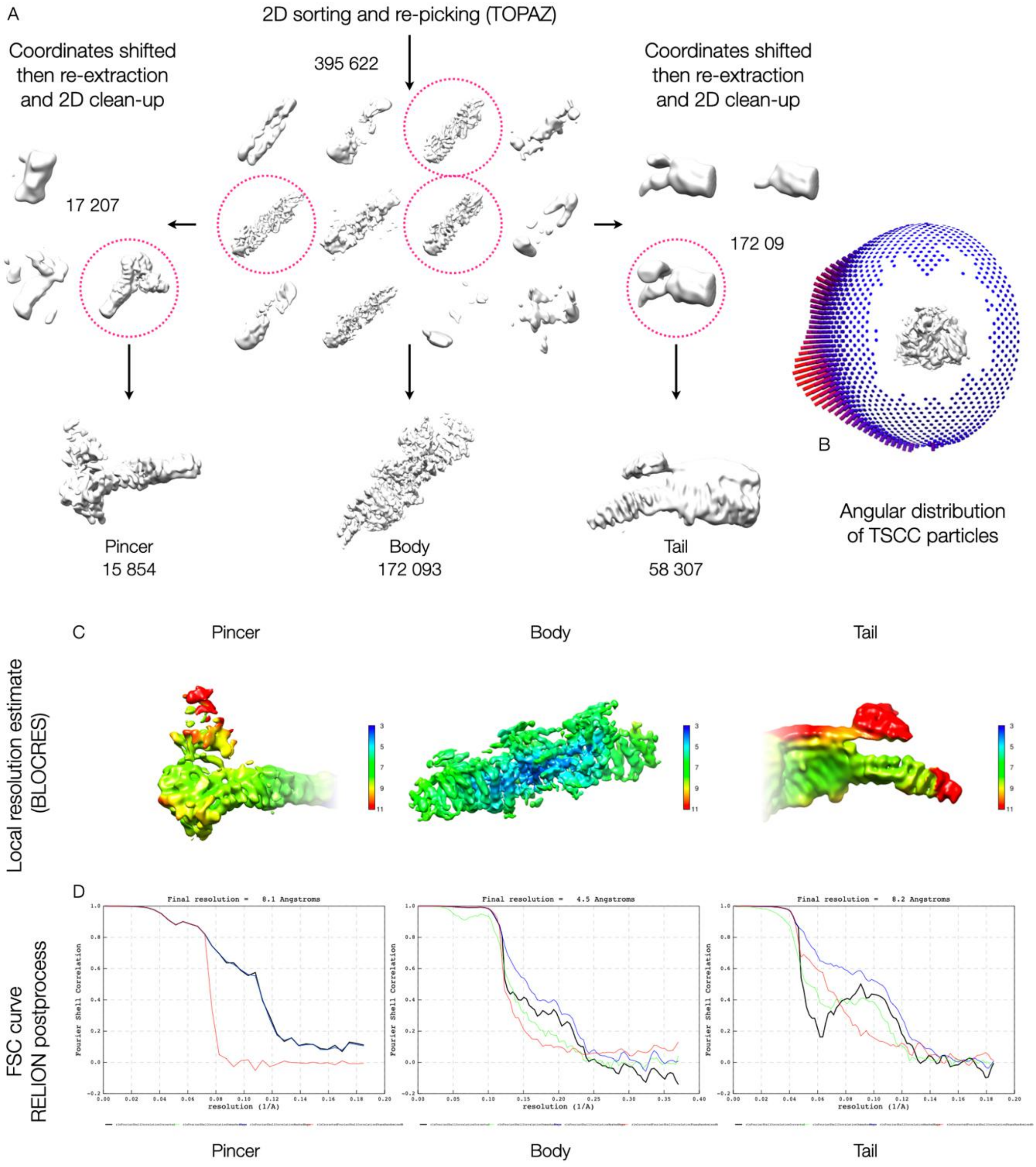


**Supplementary Figure 3. Cryo-EM data processing scheme.** A. Flow-chart showing data processing procedure (see Materials and Methods for details). Particle numbers are indicated next to each processing step. Each of the “pincer”, “body”, and “tail” regions were processed independently after isolating 2D classes representing each of these views. B. Angular distribution of TSCC particles on the grid, demonstratingpreferred orientation and absence of end-on views. C. Local resolution maps estimated for each reconstruction using BLOCRES. D. The global resolution of each final map by FSC using a cutoff of 0.143 in RELION.

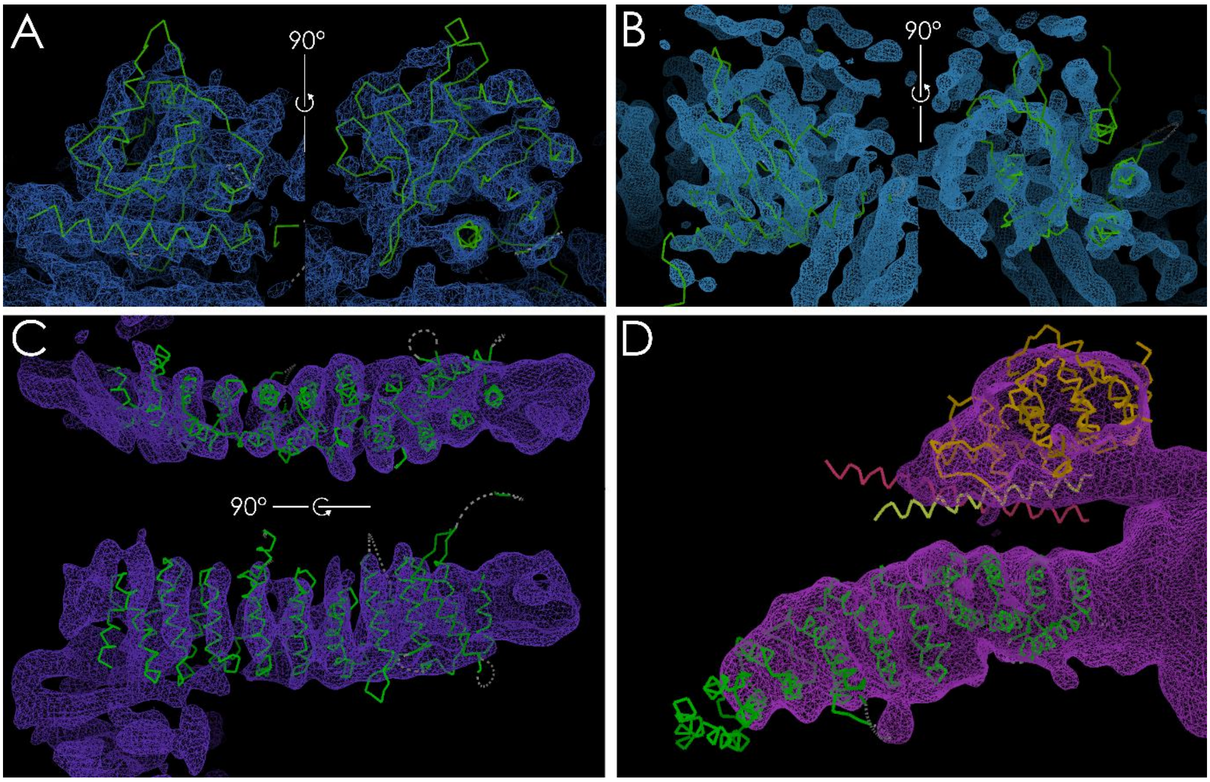


**Supplementary Figure 4. Alignment of known crystal structures in cryo-EM reconstructions.** A. The *C. thermophilum* $TSC2_{GAP}$ crystal structure (PDB ID: 6SSH; shown as backbone in green) (Hansmann *et al*, 2020) fits unambiguously into the region of density bound against the TSC2 HEAT repeat within the "body". The human GAP domain is therefore highly structurally homologous. B. The dimerisation domain of the human RapGAP crystal structure (PDB ID: 3BRW; shown as backbone in green)(Scrima *et al*, 2008) fits into the dimerisation interface between the two TSC2 HEAT repeats. The topology of the β-sheets is conserved. C. The *C. thermophilum*TSC2 N-terminal HEAT repeat fragment crystal structure (PDB ID: 5HIU; shown as backbone in green) (Zech *et al*, 2016) fits into the "pincer", indicating the α-solenoid extends from C-terminus, at the dimerisation interface, to N-terminus, at either end of the elongated complex. D. The *C. thermophilum* TSC2 N-terminal α-solenoid, as shown in C., and the TBC1D7 crystal structure (PDB ID: 5EJC) (Qin *et al*, 2016) fit into density of the "tail" and "barb". The TBC1D7 density is weak, indicating a less well-ordered interaction between the TSC1 coiled-coil-TBC1D7 junction and TSC2.

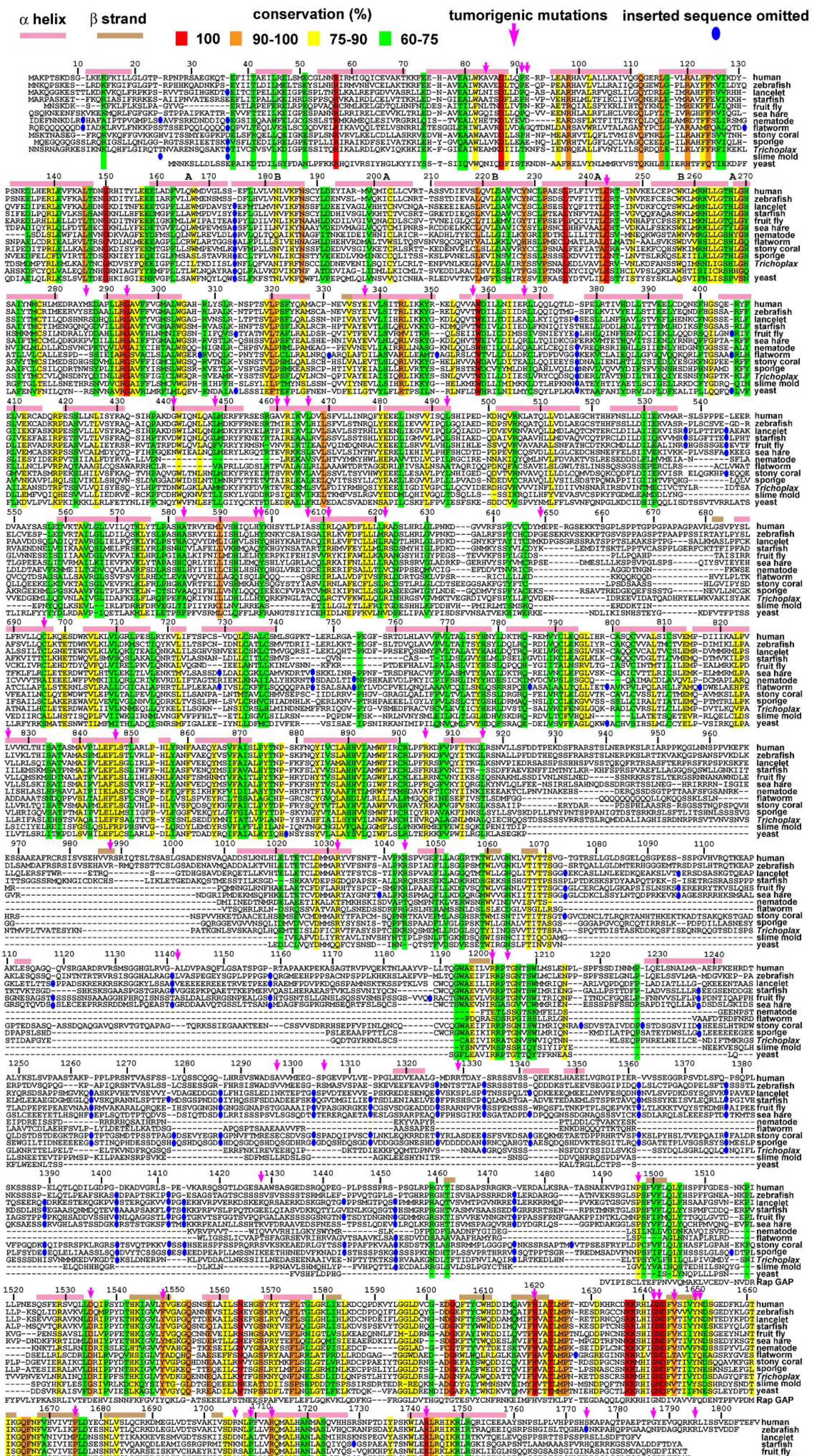

α helix
β strand
conservation (%)
100
90-100
75-90
60-75
tumorigenic mutations
inserted sequence omitted
human
zebrafish
lancelet
starfish
fruit fly
sea hare
nematode
flatworm
stony coral
sponge
Trichoplax
slime mold
yeast
Rap GAP

**Supplementary Figure 5. Multiple sequence alignment and secondary structural prediction of TSC2.** Human TSC2 displays average sequence conservation with close homologs for most of the protein except the GAP domain, which is highly conserved. Human TSC2 protein (1807 residues) was aligned with orthologues from zebrafish (*Danio rerio*, accession: XP_021331278), lancelet (*Bronchiostoma belcheri*, subphylum of Cephalochordata, accession: XP_019643564), starfish (*Acanthaster planci*, phylum of Echinodermata, accession: XP_022103490), fruit fly (*Drosophila melanogaster*, phylum of Arthropoda, accession: NP_524177), sea hare (*Aplysia californica*, phylum of Mollusca, accession: XP_012937341), nematode (*Brugia malayi*, phylum of Nematoda, accession: XP_001893474), flatworm (*Macrostomum lignano*, phylum of Platyhelminthus, accession: PAA85829), stony coral (*Orbicella faveolata*, phylum of Cnidaria, accession: XP_020601823), sponge (*Amphimedon queeslandica*, phylum of Porifera, accession: XP_019853372), *Trichoplax adhaerens* (phylum of Placozoa, accession: XP_002116811), slime mold (*Planoprotostelium fungivorum*, kingdom of Protista, accession: PRP73953), and yeast (*Schizosaccharomyces pombe*, kingdom of Fungi, accession: CAB52735), as well as human RapGAP by the Clustal W method using MegAlign in the LASERGENE software. The tumorigenic mutations of TSC2 are indicated with magenta arrows.

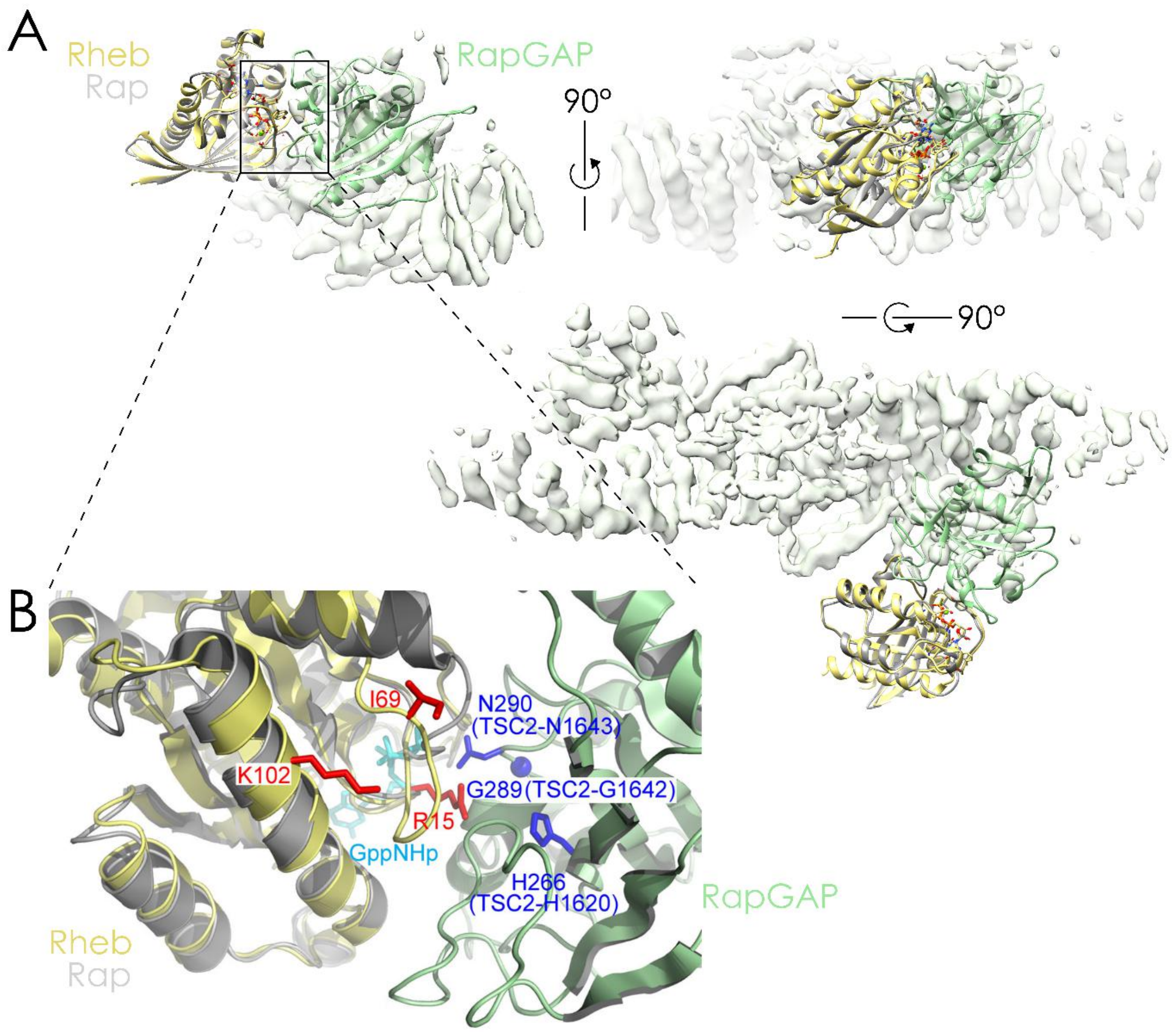


**Supplementary Figure 6. Rheb docking and TSC2-GAP-Rheb interactions based on sequence and structural homology to Rap GTPase-RapGAP.** A. The crystal structure of Rheb·GTP (Yu *et al*, 2005) (PDB ID: 1XTS) is superimposed onto the crystal structure of the Rap GTPase-RapGAP crystal structure (Scrima *et al*, 2008) (PDB ID: 3BRW). The RapGAP domain fitted into density we have assigned to the TSC2 GAP domain in our "body" reconstruction, highlighting how Rheb is captured in this conformation. Rheb is coloured yellow, the Rap GTPase grey, and the RapGAP domain light green. B. The crystal structure of Rheb is superimposed onto the crystal structure of the RapGAP-Rap complex as in A. Rap-contacting residues of RapGAP which correspond to tumorigenic mutations on TSC2 are highlighted in blue. The Rheb residues which are conserved in Rheb homologues but are different in all other Ras family GTPase members are highlighted in red. Interestingly, these residues of Rheb all point toward TSC2-GAP.

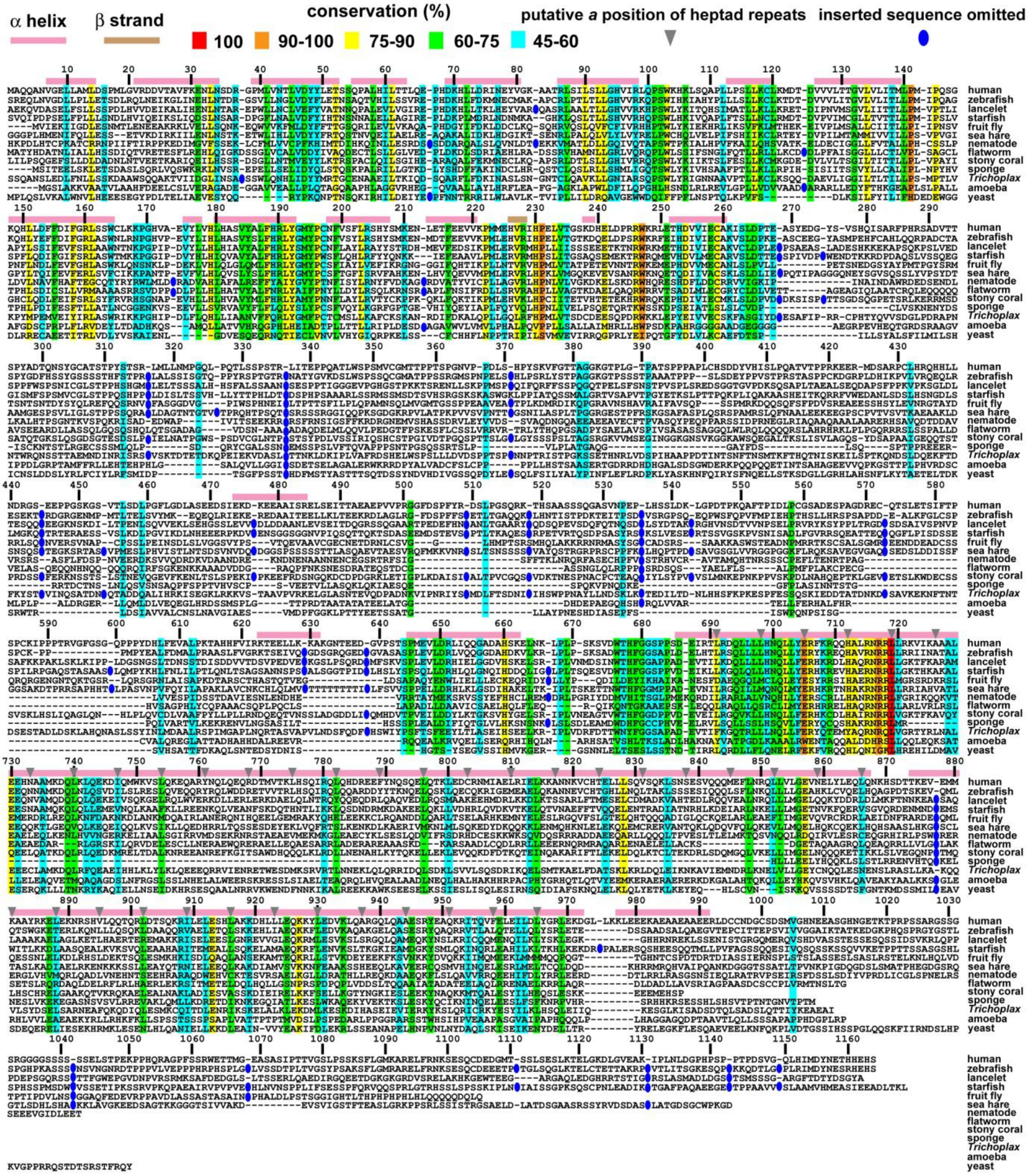


**Supplementary Figure 7. Multiple sequence alignment and secondary structural prediction of TSC1.** Human TSC1 protein (1164 residues) was aligned with orthologues from zebrafish (*Danio rerio*, accession: NP_001269321), lancelet (*Bronchiostoma belcheri*, accession: XP_019640975), starfish (*Acanthaster planci*, accession: XP_022085127), fruit fly (*Drosophila melanogaster*, accession: CCB63097), sea hare (*Aplysia californica*, accession: XP_005108980), nematode (*Ancyclostoma ceylanicum*, accession: EPB75620), flatworm (*Macrostomum lignano*, accession: PAA78603), stony coral (*Stylophora pistillata*, accession: XP_022782800), sponge (*Amphimedon queenslandica*, accession:

XP_019858430), *Trichoplax adhaerens* (accession: XP_002109434), amoeba (*Acanthamoeba castellanii*, accession: XP_004339658), and yeast (*Schizosaccharomyces pombe*, accession: CAA91078), by the Clustal W method using MegAlign in the LASERGENE software.

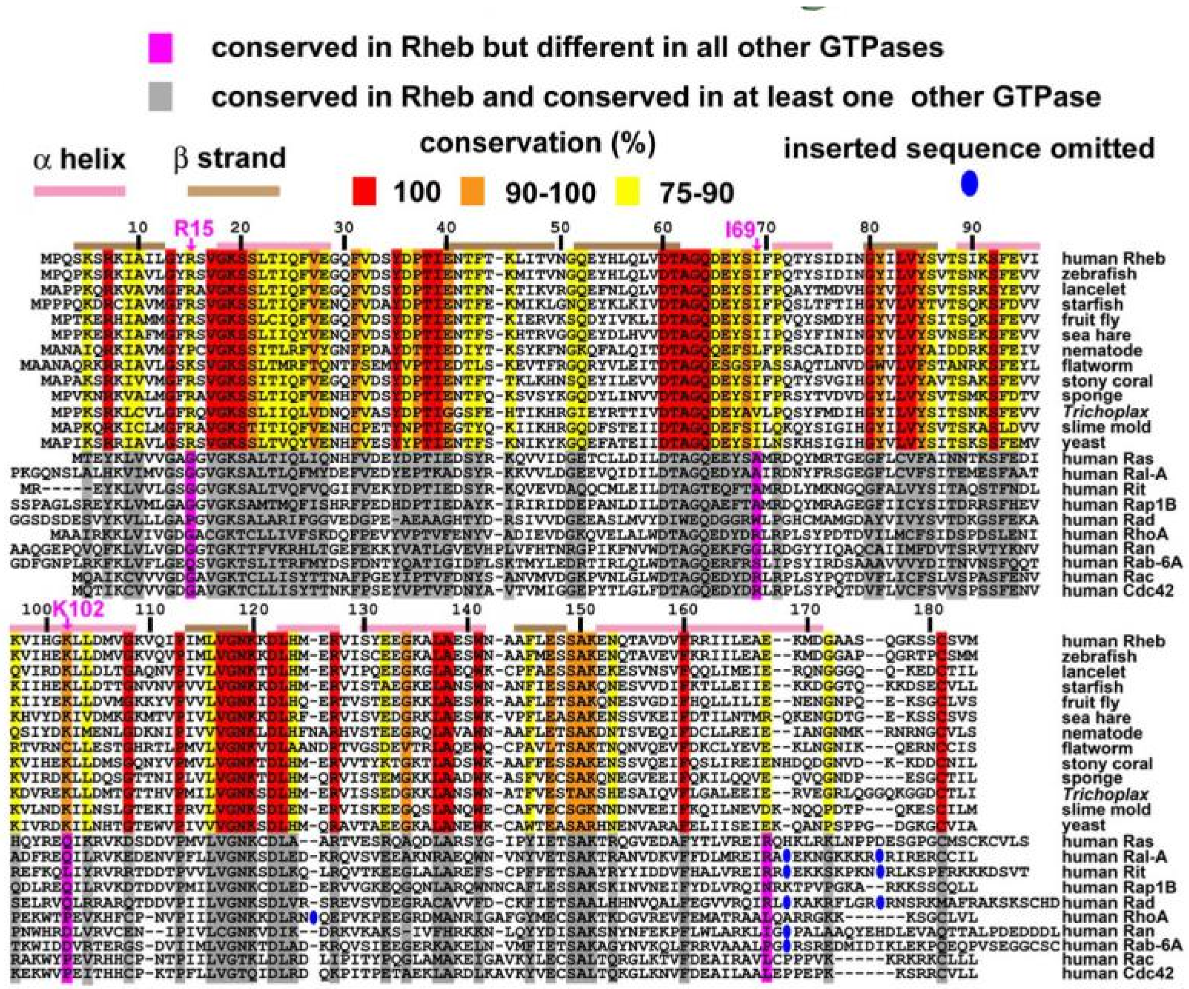


**Supplementary Figure 8. Multiple sequence alignment of Rheb and other Ras family members show that R15, I69, and K102 of Rheb are unique among Ras family.** Residues unique to Rheb, *i.e.*, those which are conserved in Rheb homologues but are different in all other Ras family GTPases are highlighted in magenta.

**Supplementary Table1. Frequent tumourigenic missense mutations of TSC2 in the tuberous sclerosis database.** Tabulated tumourigenic missense mutations of TSC2 with over five entries in the Leiden Open Variation Database:

[ http://chromium.lovd.nl/LOVD2/TSC/home.php?select_db=TSC2 ].

| residue | Tumorigenic missense mutations of TSC2 | | | | Total entries |
|---|---|---|---|---|---|
| A84 | A84V (5 entries) | | | | 5 |
| P91 | P91L (6 entries) | | | | 6 |
| E92 | E92V (8 entries) | | | | 8 |
| C244 | C244R (2 entries) | C244Y (3 entries) | | | 5 |
| M286 | M286V (10 entries) | M286T (1 entry) | | | 11 |
| G294 | G294R (1 entry) | G294E (2 entries) | G294V (2 entries) | | 5 |
| E337 | E337K (5 entries) | | | | 5 |
| A357 | A357V (5 entries) | | | | 5 |
| R367 | R367Q (17 entries) | R367P (1 entry) | | | 18 |
| G440 | G440S (7 entries) | | | | 5 |
| L448 | L448P (4 entries) | L448R (1 entry) | | | 5 |
| A460 | A460T (11 entries) | | | | 11 |
| R462 | R462G (2 entries) | R462C (1 entry) | R462H (6 entries) | R462P (2 entries) | 11 |
| L466 | L466P (4 entries) | L466R (1 entry) | | | 5 |
| L493 | L493V (4 entries) | L493P (2 entries) | | | 6 |
| A583 | A583T (8 entries) | | | | 8 |
| H597 | H597Y (3 entries) | H597P (1 entry) | H597R (7 entries) | H597L (1 entry) | 12 |
| Y598 | Y598H (5 entries) | Y598L (2 entries) | | | 7 |
| A607 | A607T (7 entries) | A607S (1 entry) | A607E (1 entry) | | 9 |
| R611 | R611W (44 entries) | R611G (4 entries) | R611Q (87 entries) | R611P (1 entry) | 136 |
| R622 | R622W (10 entries) | R622Q (1 entry) | R622P (4 entries) | | 15 |
| M649 | M649V (1 entry) | M649L (1 entry) | M649T (6 entries) | | 8 |
| C696 | C696R (4 entries) | C696Y (2 entries) | | | 6 |
| L826 | L826M (8 entries) | L826P (3 entries) | | | 11 |
| L847 | L847P (5 entries) | L847R (1 entry) | | | 6 |
| R905 | R905W (49 entries) | R905G (2 entries) | R905Q (17 entries) | | 68 |
| L916 | L916P (3 entries) | L916R (2 entries) | | | 5 |
| R988 | R988C (1 entry) | R988H (1 entry) | R988P (3 entries) | | 5 |
| R1032 | R1032P (8 entries) | | | | 8 |
| R1044 | R1044K (2 entries) | R1044T (1 entry) | R1044M (2 entries) | | 5 |
| A1141 | A1141T (1 entry) | A1141V (11 entries) | | | 12 |

| | | | | | | |
|---|---|---|---|---|---|---|
| R1200 | R1200W (28 entries) | R1200Q (1 entry) | R1200P (3 entries) | | | 32 |
| T1203 | T1203P (4 entries) | T1203K (1 entry) | | | | 5 |
| A1297 | A1297T (8 entries) | | | | | 8 |
| P1305 | P1305S (1 entry) | P1305L (5 entries) | | | | 6 |
| R1329 | R1329H (12 entries) | | | | | 12 |
| A1429 | A1429S (10 entries) | | | | | 10 |
| P1497 | P1497T (3 entries) | P1497S (2 entries) | P1497R (3 entries) | P1497L (4 entries) | | 12 |
| D1535 | D1535Y (2 entries) | D1535A (2 entries) | D1535V (1 entry) | | | 5 |
| Y1549 | Y1549N (1 entry) | Y1549H (1 entry) | Y1549C (4 entries) | | | 6 |
| H1620 | H1620Y (5 entries) | H1620R (1 entry) | | | | 6 |
| G1642 | G1642D (5 entries) | | | | | 5 |
| N1643 | N1643H (3 entries) | N1643S (2 entries) | N1643I (1 entry) | N1643K (1 entry) | | 7 |
| V1646 | V1646M (3 entries) | V1646L (2 entries) | V1646G (1 entry) | | | 6 |
| Y1650 | Y1650N (2 entries) | Y1650H (2 entries) | Y1650S (1 entry) | Y1650C (4 entries) | | 9 |
| N1651 | N1651H (2 entries) | N1651S (14 entries) | | | | 16 |
| P1675 | P1675L (55 entries) | P1675Q (2 entries) | P1675R (4 entries) | | | 61 |
| R1706 | R1706C (3 entries) | R1706H (2 entries) | | | | 5 |
| P1709 | P1709L (12 entries) | P1709R (2 entries) | | | | 14 |
| R1713 | R1713H (9 entries) | R1713P (3 entries) | | | | 12 |
| R1743 | R1743G (2 entries) | R1743W (47 entries) | R1743Q (49 entries) | R1743P (6 entries) | R1743L (2 entries) | 106 |
| S1774 | S1774T (9 entries) | | | | | 9 |
| G1787 | G1787S (6 entries) | | | | | 6 |
| R1795 | R1795C (7 entries) | R1795L (1 entry) | | | | 8 |
